# A dual diffusion model enables 3D binding bioactive molecule generation and lead optimization given target pockets

**DOI:** 10.1101/2023.01.28.526011

**Authors:** Lei Huang, Tingyang Xu, Yang Yu, Peilin Zhao, Ka-Chun Wong, Hengtong Zhang

## Abstract

Structure-based generative chemistry aims to explore much bigger chemical space to design a ligand with high binding affinity to the target proteins; it is a critical step in *de novo* computer-aided drug discovery. Traditional *in silico* methods suffer from calculation inefficiency and the performances of existing machine learning methods could be bottlenecked by the auto-regressive sampling strategy. To address these concerns, we herein have developed a novel conditional deep generative model, PMDM, for 3D molecule generation fitting specified target proteins. PMDM incorporates a dual equivariant diffusion model framework to leverage the local and global molecular dynamics to generate 3D molecules in a one-shot fashion. By considering the conditioned protein semantic information and spatial information, PMDM is able to generate chemically and conformationally valid molecules which suitably fit pocket holes. We have conducted comprehensive experiments to demonstrate that PMDM can generate drug-like, synthesis-accessible, novel, and high-binding affinity molecules targeting specific proteins, outperforming the state-of-the-art (SOTA) models in terms of multiple evaluation metrics. In addition, we perform chemical space analysis for generated molecules and lead compound optimization for SARS-CoV-2 main protease (M_pro_) by only utilizing three atoms as the seed fragment. The experimental results implicate that the structures of generated molecules are rational compared to the reference molecules, and PMDM can generate massive bioactive molecules highly binding to the targeted proteins which are not included in the training set.

## Main

Structure-based drug discovery is a critical task for modern drug development^12^ and catalysis^3^. Given a specific target protein, it aims to find proper drug molecules that bind well with the target protein. Traditional *in silicon* methods such as virtual screening discover molecules by iteratively (1) placing molecules from existing databases into the pocket cavity; and (2) filtering the molecules by the rules such as energy estimation^4^ and toxicity followed experiment essays. Despite their wide applications, such approaches suffer from two major limitations^5,6^. First, naive exhaustive searches in the massive chemical space (range from 10^60^ to 100^100^ depending on the size of desired molecules)^7^ are prohibitively costly. Second, such a workflow is limited by historical knowledge, thus infeasible to explore and generate novel molecular structures which are not recorded in the existing databases.

Fortunately, the emergence of deep learning methods has paved the way for efficient and accurate drug molecular structure learning and has greatly facilitated the exploration of chemical spaces rich in structured biological data distributions in recent years. By adopting generative models, current machine learning methods^8,9,10,11,12,13^ start from learning the underlying distribution of molecules and yield candidate molecules from perturbed hidden information. Nonetheless, these methods represent the molecules as SMILES strings (1D) or graphs (2D), but ignore the 3D-spatial information that is crucial to determine the properties of molecules. For example, a molecular graph is capable of forming various conformations with different properties in 3D space due to intramolecular interactions or the orientation of structural motifs^14^. Later, some methods^15,16,17,18^ generate 3D molecules by considering 3D-spatial information. However, these methods do not involve the pocket information to generate molecules with high binding affinity. As a result, the resulting molecules cannot fit into specific protein pockets, which is not suitable for wet experiments. This gives rise to the idea of structure-based generative chemistry, where the molecules with high binding affinity in the protein pocket are distilled. Here, the models perceive the 3D structure of the target pocket as the conditional information and the interaction between molecules and proteins to learn the conditioned density of desired molecular data.

Most recently, some generative models that enable 3D sampling of molecules within the pocket cavity have been pro-posed^19,20,21,22^. Early attempts^20^ incorporate conditioned VAE to handle the voxelized atomic density images and obtain molecules from the images by post-processing algorithm. However, it compresses the pocket structure information and fails to generate accurate molecules with fine-grained positions. Followed studies^21,22^ adopt auto-regressive frameworks to sample one atom at each time step, but are inefficient in the generation process due to the large candidate chemical space. The sequence nature of these methods leads to deviation accumulations, especially when invalid structures are generated in the early steps. Furthermore, the sequential generation process may result in the loss of global context information since molecular structures are disordered. Therefore, achieving 3D sampling of molecules within pocket cavities remains challenging.

Recently, diffusion models^23^ have received a huge amount of attention in computer vision tasks^24,25,26^, especially in point could generation^27,28,29^ which is similar to 3D molecule generation. These methods can inpaint the 3D objects by learning the joint distribution. Inspired by the success of diffusion models in computer vision tasks, we propose a one-shot generation framework named Pocket based Molecular Diffusion Model (PMDM) to tackle such issues. Figure 1 outlines the overview of PMDM. Specifically, molecular atoms with fixed pocket information are regarded as 3D point clouds and diffused in the forward process which is similar to the phenomenons in nonequilibrium thermodynamics. The goal of PMDM is to learn how to reverse such process to model a conditioned data distribution. This allows us to efficiently generate accurate molecules with high binding affinity once the pocket information is fixed. However, regular methods for 3D point clouds cannot involve edge information like chemical bond information if we represent 3D molecular geometries as 3D point clouds. Thus, we define a dual diffusion strategy to build two kinds of virtual edges. In detail, pairs of atoms with interatomic distances below a certain threshold are bonded via covalent localized edges because chemical bonds can dominate interatomic forces when two atoms are close enough to each other while global edges are linked to the remaining pairs of atoms to simulate the van der Waals force. Besides, we design an equivariant dynamic kernel that obeys the translation, rotation, reflection, and permutation equivariance of molecular geometry systems. The experiments on synthetic CrosseDocked dataset^30^ demonstrate that PMDM can generate drug-like, synthesis-accessible, novel molecules with high binding affinity against specific proteins and outperform the state-of-the-art (SOTA) models on multiple evaluation metrics.

**Figure 1.**
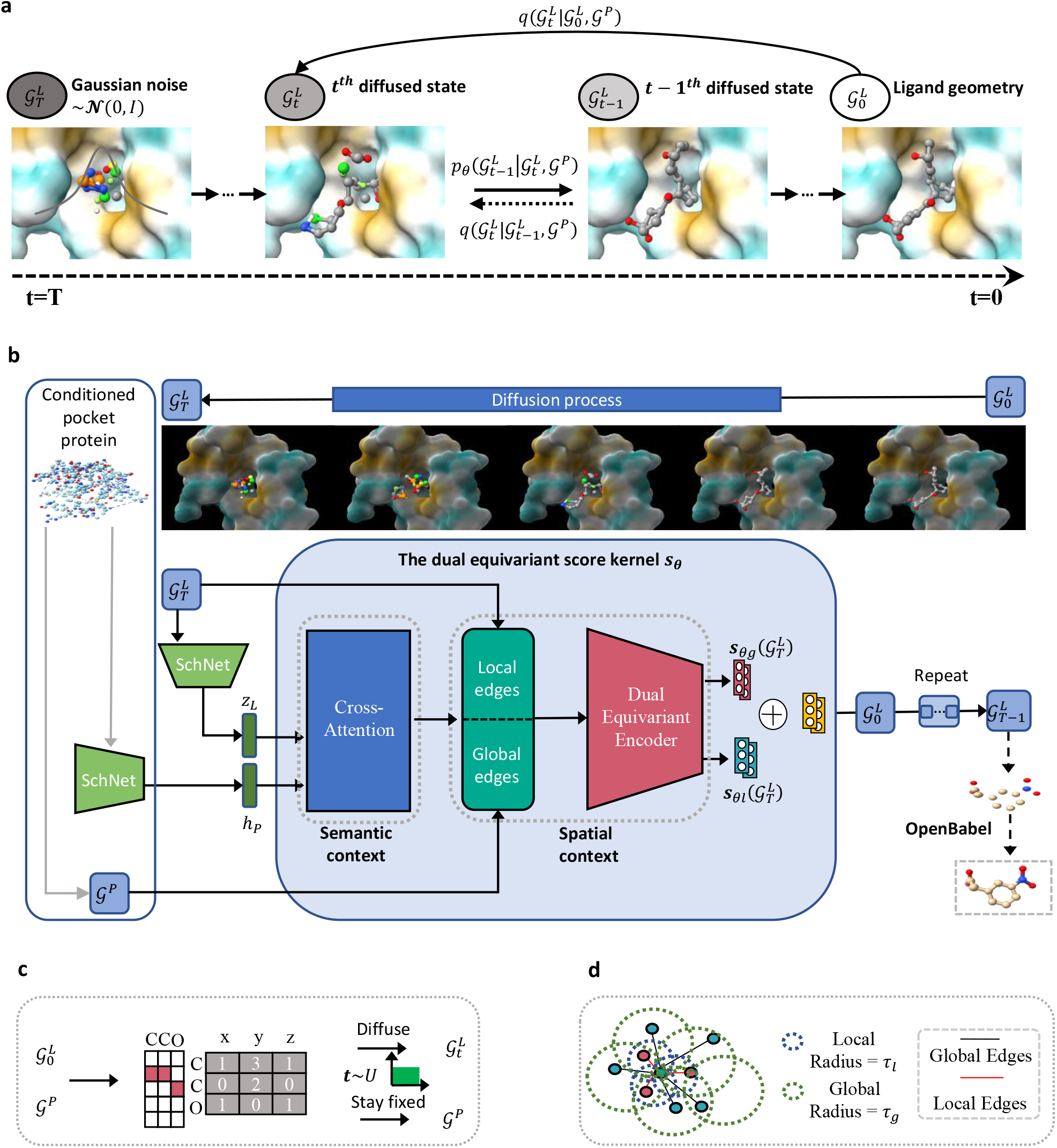
Overview of the PMDM framework. a. The diagram of diffusion process in PMDM. PMDM is based on the diffusion model, which defines two Markov processes: diffusion process and reverse process. The diffusion process iteratively adds Gaussian noises to the ligand data 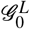 according to a variance preserve schedule while the reverse process generates a realistic ligand from the corruption state 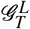 through eliminating the noise. In the training phase, any immediate state 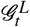 can be calculated by 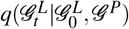, we will elaborate this desired property in section Methods. Since the diffusion process is fixed, PMDM is trained to learn the reverse probability transition distribution 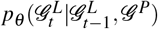. b. The structure of PMDM. PMDM is designed to generate the ligand given the target pocket protein. PMDM could encode the protein semantic context information and spatial context information. The protein point cloud data is fed into an invariant encoder SchNet to obtain the semantic representation *h*_*p*_. Then the semantic information is fused with the ligand data by the cross-attention layers. We define local and global edges for ligand point cloud data. Then the ligand data with two kinds of edges and pocket protein data go through the dual equivariant encoder which handles different edges and keeps the protein spatial information fixed to obtain the score *s*_*θ*_. This process will repeat *T* times until we obtain the realistic ligand geometry 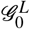, and we use OpenBabel to construct the bonds. c. The ligand and protein are represented by one-hot encoded atom types and 3D coordinates. The ligand data will be diffused to 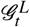 at an arbitrary time step while the protein will stay fixed during training. d. The construction of local and global edges.

## Related work

We have witnessed a variety of deep generative models have been proposed to generate molecules. A plethora of studies consider generating molecules via advanced generative methods, including variational autoencoder (VAE)-based models^31^, generative adversarial network (GAN)^32^, normalizing flows^9,12,17^ and diffusion models^18,16^. These methods, however, do not incorporate pocket information, which are far away from the clinical application. Then, the SBDD task which depends on the binding pocket is gaining attention from researchers. Early studies focus on 1D or 2D molecule generation based on the pocket structure. Skalic et al.^33^ propose a framework which is based on a variant GAN to generate SMILES strings of ligands after encoding the molecules strings in the shared space with the pocket protein. Xu et al.^34^ employ the conditional RNN to train two descriptors which contain the 3D information of pocket to generate compounds. However, these methods can only generate molecules in SMILES sequence format (1D) or graph (2D) which cannot verify the fitness of the target pocket although they considered the 3D information of the pocket.

Recently, research on generating molecules in 3D space has been proposed for accommodating the needs of fitting the specific protein structure. Masuda et al.^20^ firstly incorporate conditional VAE to generate atomic densities of the desired ligand based on the pocket protein. They use convolutional neural networks to encode the density girds into separate ligand and protein latent spaces. Compared with previous work which can only generate small molecules, this method can generate more drug-like 3D molecules. However, this method does not consider the equivariance of molecular geometry. Besides, it is hard to scale to large proteins due to the voxel design. To tackle this issue, Luo et al.^21^ model the atom probability by graph neural networks and equip the mask-fill schema to estimate the landscape of the pocket. Liu et al.^22^ further incorporate distance and angles embeddings to place the atom one by one. The existing generative models typically adopt the auto-regressive strategy to sample the atom sequentially, which enables the current atom to learn the historically placed atom information. Nonetheless, these methods have inherent limitations: (1) the models may suffer from deviation accumulations especially when invalid structures are generated in the early steps; (2) the sequential sampling algorithm that relies on MCMC does not consider the global context information; (3) auto-regressive models place one atom at a time, thus the number of the sequential sampling is the same as the length of the ligand, making it time-consuming to generate large-scale molecules.

## Materials

### Data

We conduct experiments to evaluate the generative performance of PMDM on the CrossDocked dataset^30^. This dataset contains 22.5 million docked protein-ligand pairs and each pair has different poses to multiple pockets across the Protein Data Bank. For a fair comparison, we follow previous work^21^ to only choose the binding pose data whose root-mean-squared deviations (RMSD) is less than 1Å. The dataset is then refined through clustering at 30% sequence identity using MMseqs2^35^, finally we obtain 100,000 pairs for training and 100 pairs for evaluation.

### Metrics

We adopt widely-used metrics^36,21^ to evaluate the quality of molecules generated by PMDM: (1) **Vina Score** estimates the binding affinity between the ligand and the target pocket which is the most important measurement to evaluate how the generated molecule fits into the protein pocket of interest; (2) **High affinity** is the percentage of the molecules whose **Vina Score** is higher than the ground truth molecule in the test set; (3) **QED** estimates the drug-likeness of the molecule via combining several desirable molecular properties; (4) **SA** (synthetic accessibility) measures the molecule synthetic accessibility; (5) **Lipinski** measures how many rules the drug follows five Lipinski’s rules^37^; (6) **LogP** indicates the octanol-water partition coefficient, which should be between -0.4 and 5.6 if the molecule is a good drug candidate^38^; (7) **Diversity** represents the average pairwise Tanimoto dissimilarity of the generated molecules targeting for each pocket. (8) **Time** is the average time to generate 100 samples for each pocket across all the targets.

### Baseline models

We compared PMDM with SOTA models for the SBDD task including CVAE^20^ and AR-SBDD^21^. Both of the methods adopt an auto-regressive strategy to generate samples. Besides, we also report the calculation results of molecules in the test set for a more comprehensive comparison.

## Results

### Evaluation of PMDM on the general metrics

We generate 100 molecules for each target protein in the test set (10000 molecules in total). Here, the size of the generated molecules is sampled from the size distribution of the training set. The overall results of PMDM and the baseline models are presented in Table 1. We observe that PMDM outperforms all the baseline models on almost metrics except SA and Diversity. According to the Vina score, PMDM is able to generate the molecules with high affinity to the pocket (−7.472 ± 2.90) which is 20.2% better than the best baseline, AR-SBDD. Besides, PMDM surpasses AR-SBDD on QED (0.619 ± 0.13) by 23.3 %, and Lipinski (4.991 ± 0.09) by 4.2 %. The logP value of PMDM within the compliance range (−0.4 5.6) implies that the molecules generated by PMDM hold greater promise as drug candidates, which is crucial for clinical trials. For the SA, PMDM achieves competitive results compared to AR-SBDD and performs better than CVAE. On the other hand, the diversity of generated molecules should fall within a reasonable range so that the ability to explore the molecular space confined by protein pockets is high enough to discover novel molecules. As in Table 1, the diversity of PMDM is lower than that of AR-SBDD but higher than that of CVAE, implying that our model satisfies this desired property.

**Table 1.** The comparison of 10000 generated molecules of PMDM and baseline models on the CrossDocked dataset. We generate 100 samples for each target pocket.

| Methods | Vina Score (kcal/mol) ↓ | High Affinity ↑ | QED ↑ | SA ↑ | Lipinski ↑ | LogP | Diversity | Time (seconds) ↓ |
| --- | --- | --- | --- | --- | --- | --- | --- | --- |
| CVAE | $-6.144 \pm 1.57$ | 0.238 | $0.369 \pm 0.22$ | $0.590 \pm 0.15$ | $4.367 \pm 1.14$ | $-0.140 \pm 2.73$ | $0.654 \pm 0.12$ | - |
| AR-SBDD | $-6.215 \pm 1.54$ | 0.267 | $0.502 \pm 0.17$ | **$0.675 \pm 0.14$** | $4.787 \pm 0.50$ | $0.257 \pm 2.01$ | $0.742 \pm 0.09$ | $19659 \pm 14704$ |
| **PMDM** | **$-7.572 \pm 2.50$** | **0.628** | **$0.594 \pm 0.12$** | $0.611 \pm 0.16$ | **$4.975 \pm 0.16$** | $0.301 \pm 1.01$ | $0.709 \pm 0.10$ | **906 ± 1097** |
| Test set | -7.024 | - | 0.466 | 0.725 | 1.413 | 0.929 | - | - |

Notably, the molecules generated by PMDM perform even better than those in the test set on Vina Score, QED, and Lipinski, suggesting that PMDM has great potential to generate more drug-like molecules with higher affinity outside the distribution of the dataset. The one-shot nature of PMDM ensures that the model effectively considers the global information of the molecule, rather than sampling the local optimum atom like auto-regressive methods, which is time-consuming. Thus, as a one-shot method, PMDM is able to sample molecules up to two times faster than auto-regressive models while achieving better or competitive performance.

### Analysis of PMDM on local geometries

Although conventional metrics can reflect the quality of generated molecules to a certain extent, the quality of the sub-structures of generated molecules also needs to be considered when evaluating model performance. We select several pocket proteins to visualize as representative samples for sub-structure analysis. As depicted in Figure 2, we choose 14GS, 2RMA, and 3AF2 as the targeted pocket protein. We observe that the AR-SBDD tends to generate the three-atom rings while our proposed model PMDM avoids generating such unstable rings. Although the dataset contains only 3% three-atom rings, AR methods generate more of these unstable structures, which means that these methods get stuck in local optima and fail to learn the data distribution well. Instead, PMDM can consider the shape of the pocket hole and generate larger and more complicated rings which are shown in the 3AF2 pocket samples.

**Figure 2.**
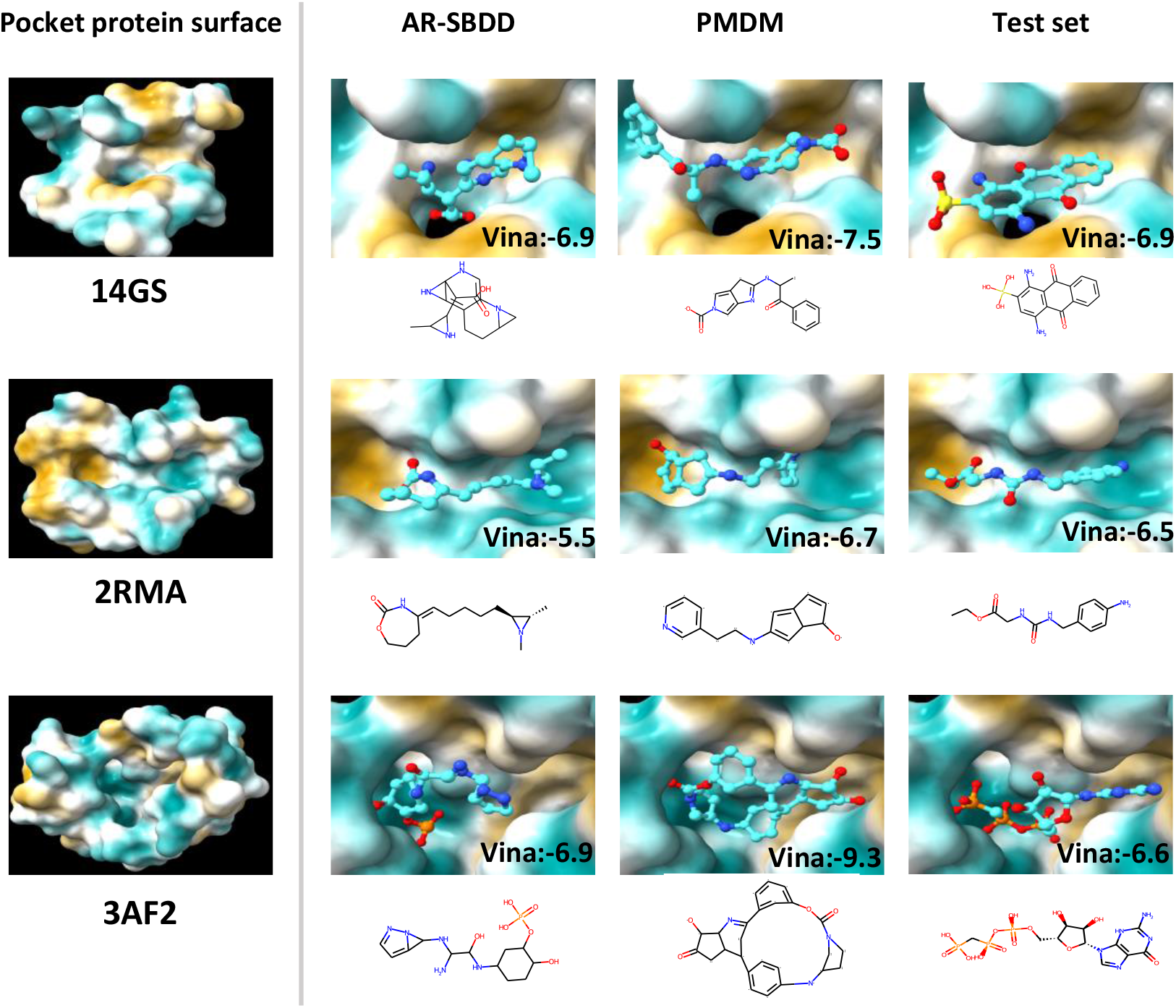
The comparison of the example molecules which are generated by AR-SBDD and PMDM, and from the test set, respectively. The molecules are targeting GLUTATHIONE S-TRANSFERASE (PDB id: 14GS), a complexed Crystal Structure of Cyclophilin (PDB id: 2RMA), and Pantothenate kinase (PDB id: 3AF2), respectively.

To further quantify the ring sub-structure of the molecules generated by those methods, we report the proportion of molecules containing rings of different sizes in the training set, the test set, and the generated sets from the methods. As presented in Figure 3b, the molecules generated by PMDM contain few unstable rings, including the three-atom ring and the four-atom ring. In contrast, both auto-regressive methods are prone to generate a relatively large amount of unstable molecules with three-atom rings and four-atom rings. Besides, PMDM is inclined to generate more molecules with five-atom rings and six-atom rings, where hydrogen bonds occur most frequently. Such sub-structures are actively used in drug design. Another meaningful phenomenon is that PMDM generates molecules in a similar proportion to the test set, which indicates that PMDM can learn the data distribution without bias.

**Figure 3.**
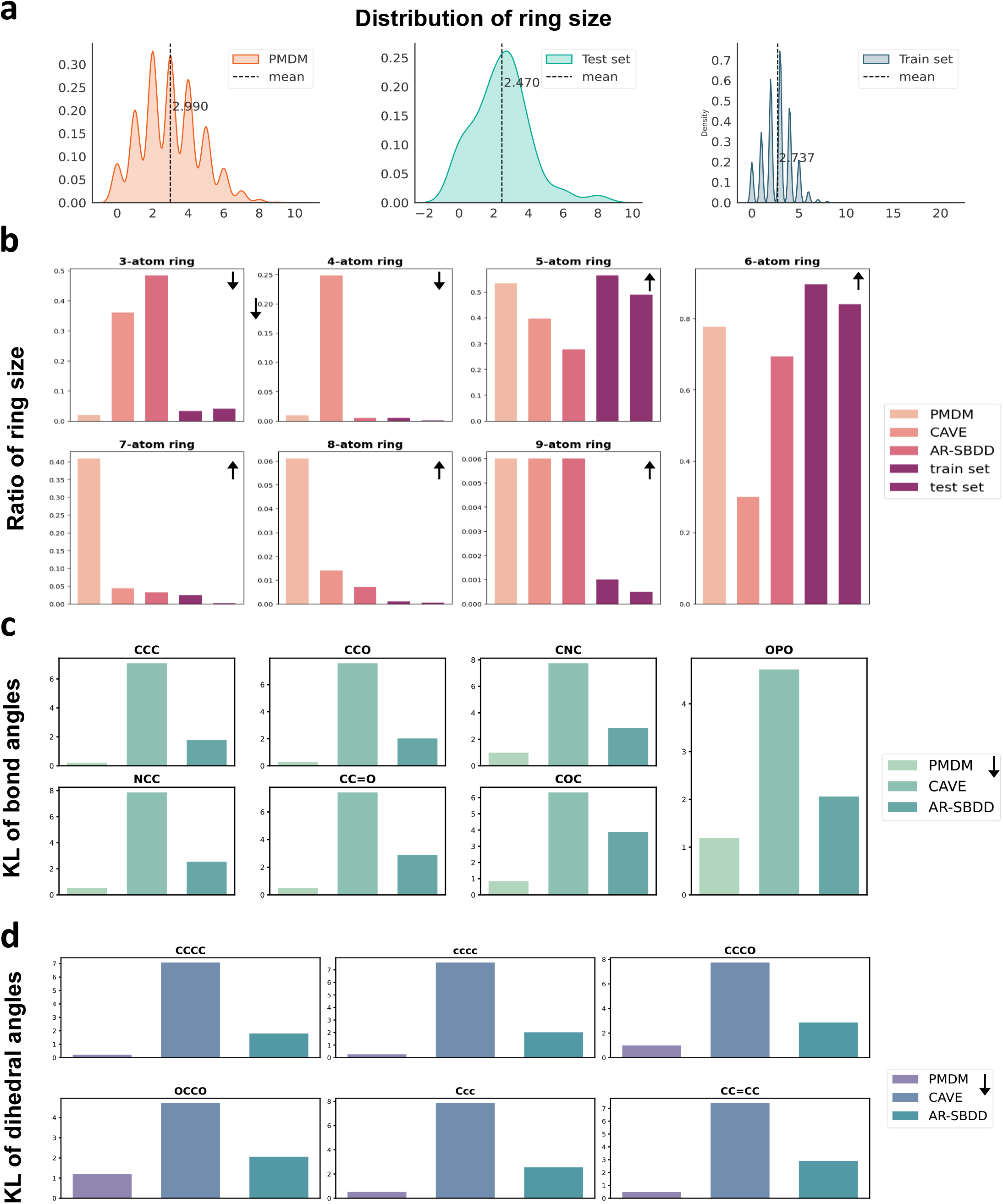
Local geometry analysis. a.The distribution of the number of rings of molecules generated by PMDM. b. The ratio of the molecules which contain rings of different sizes. c. The KL divergence of the bond angles of generated molecules from models with the test set. d. The KL divergence of the dihedral angles of generated molecules from models with the test set.

Figure 3 presents the ring number distribution of molecules generated by PMDM and the molecules in the test set and the train set. The distribution of the PMDM is close to the test set and the train set. The molecules generated by PMDM contain around 2.990 rings while the molecules in the test set and the train set contain 2.470 rings and 2.737 rings on average. Overall, the results suggest that PMDM is able to learn the ring sub-structure size distribution from a local perspective and the distribution of ring numbers from a global perspective.

We also screen out the common bond pairs and triples according to previous work^39^ and then adopt RDKit to calculate the bond angles and the dihedral angles in radians. We measure the distribution of the bond angles and dihedral angles of the generated molecules and reference molecules and then assess the distribution deviation by utilizing the Kullback-Leibler (KL) divergence. As reported in 3c and 3d, the molecules generated by PMDM demonstrate the lowest KL divergence in all the bond pair patterns and bond triple patterns among all the models. The results indicate that the PMDM is capable of capturing the local atom geometry of the data.

### Analysis of PMDM on chemical space distribution

Having analyzed the local geometry of molecules generated by PMDM, we then evaluate the generated molecular chemical space distribution from a global perspective. Since the three-dimensionality of the chemical structures is the essence of molecular design in medicinal chemistry, we also place our focus on the shape of chemical structures. Herein we adopt 2D and 3D molecular fingerprints including Morgan^40^, RDKit and USRCAT (Ultrafast Shape Recognition with CREDO Atom Types)^41^ fingerprints to represent the chemical space of generated molecules and test set molecules. Specifically, we utilize the Extended-Connectivity Fingerprints(ECFP) which are based on the Morgan algorithm to assign the unique identifiers after preset iterations. This kind of fingerprint takes atom types including connectivity and chemical features such as Donor and Acceptor and the neighbourhood of each atom into account. RDKit fingerprint is designed to measure the molecular 2D substructure by considering the atom types and bond types, which is inspired by Daylight fingerprint. In contrast, USRCAT improved USR(Ultrafast Shape Recognition) algorithm by incorporating pharmacophoric information to measure the molecular 3D shape. The visualization of the chemical space distribution using t-SNE is presented in Figure 4. The chemical space of molecules generated by PMDM can cover the molecules from the test set in the 2D substructure space, indicating that PMDM can correctly model the 2D chemical space of the training set (Figure 4a and Figure 4b). As shown in Figure 4c, the 3D chemical space of the generated molecules can basically capture the space of the test molecules due to the complexity of the conformations. Despite the incomplete coverage of the reference chemical space, there are no significant distribution mismatches between the generated and test set molecules.

**Figure 4.**
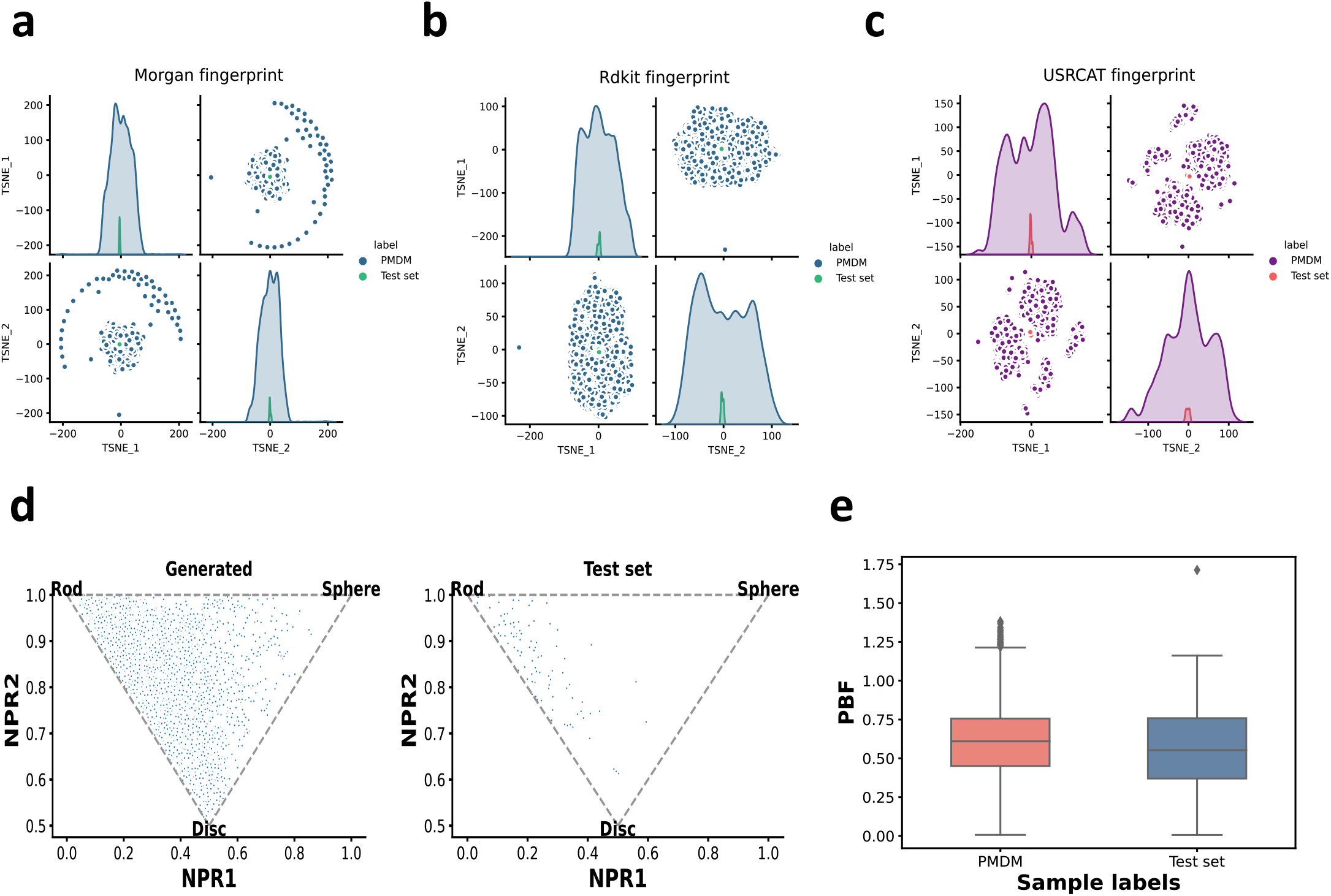
The chemical space distribution visualization. a. Morgan. b. RDKit. c. USRCAT fingerprints using t-SNE in two dimension space. 3D chemical structure measured by chemical descriptors. d. Shape distribution of generated (left) and test set (right) molecules, which is visualized using the Normalized Principal Moment of Inertia ratios(NPR) descriptors. e. The Plane of Best Fit (PBF) descriptor values.

Since the shape of 3D chemical structures is crucial for evoking molecular recognition activities with biological targets^42^, we consider leveraging molecular descriptors to characterize the three-dimensionality of molecular structures beyond the aforementioned fingerprints. Here, we adopt two widely adopted molecular descriptors: Principal Moments of Inertia (PMI)^43^ and Plane of Best Fit (PBF)^44^, to investigate the specific 3D shapes from two perspectives. Specifically, the PMI descriptors can reflect the extent to which a given molecular geometry is rod-shaped, disc-shaped or sphere-shaped while the PBF descriptors introduce the plane of best fit across all the heavy atoms of a molecule with a given conformation and calculate the distance of heavy atoms from the plane. Figure 4d depicts the Normalized Principal Moment of Inertia ratios(NPR) on a ternary plot. The closer a point is to the three corners, the more its morphology exhibits these primitive shape classes. We can observe that the generated molecules exhibit a similar gather tend to the molecules from the test set. Both the generated and test set molecules are prone to gather around the rod corner of the triangle. Furthermore, the generated molecules even touch the disc corner and sphere corner which are not covered by the original test data distribution, indicating that PMDM can not only learn the molecule 3D shape distribution of the dataset but also can explore novel shapes beyond the dataset by importing the random information, which can alleviate the out of distribution (OOD) problems in machine learning. As shown in Figure 4g, the PBF values of generated molecules and test set molecules show suitable matches, which means the degree of distance from the 2D shape is similar. To summarize, PMDM can correctly model the distribution of important 3D and 2D molecular structures and has the potential to guide a more comprehensive exploration to develop novel drug-like structures.

### Lead optimization

To further investigate the practical implications of PMDM, we apply the trained model to generate the molecules targeted for SARS-CoV-2 related proteins with high affinities. Herein, we select SARS-CoV-2 main protease (M^pro^) as a test case to perform *de novo* noncovalent inhibitors design following the previous work^45^. M^pro^ in SARS-CoV-2 is the main protease which can cleave the polyproteins at multiple positions to cleave the polyproteins, enabling it to be treated as a viable drug target. Recently, Zhang et al.^46^ redesigned the weak hit perampanel to develop a series of potent noncovalent and nonpeptidic inhibitors targeting M^pro^. In contrast to the peptide-like molecules that covalently bind to the residue (Cys145), the designed inhibitors avoid the issues of proteolytic degradation, limited antiviral activity, and molecular promiscuity toxicities. Figure 5a shows the crystal structure of one of the inhibitors complexed with M^pro^ with high bioactivity which is included in the Protein Data Bank (PDB ID: 7L11). There are several features which contribute to the high binding affinity of the molecule with M^pro^, (1) the four rings of the molecule are being placed in the four sites (S1^*′*^, S2, and S3) of the pocket; (2) the carbonyl group in the central pyridinone ring forms a hydrogen bond with the backbone NH of residue Glu166; (3) the Nitrogen in the pyridinone ring connected by the central pyridinone ring forms a hydrogen bond with the residue His 163.

**Figure 5.**
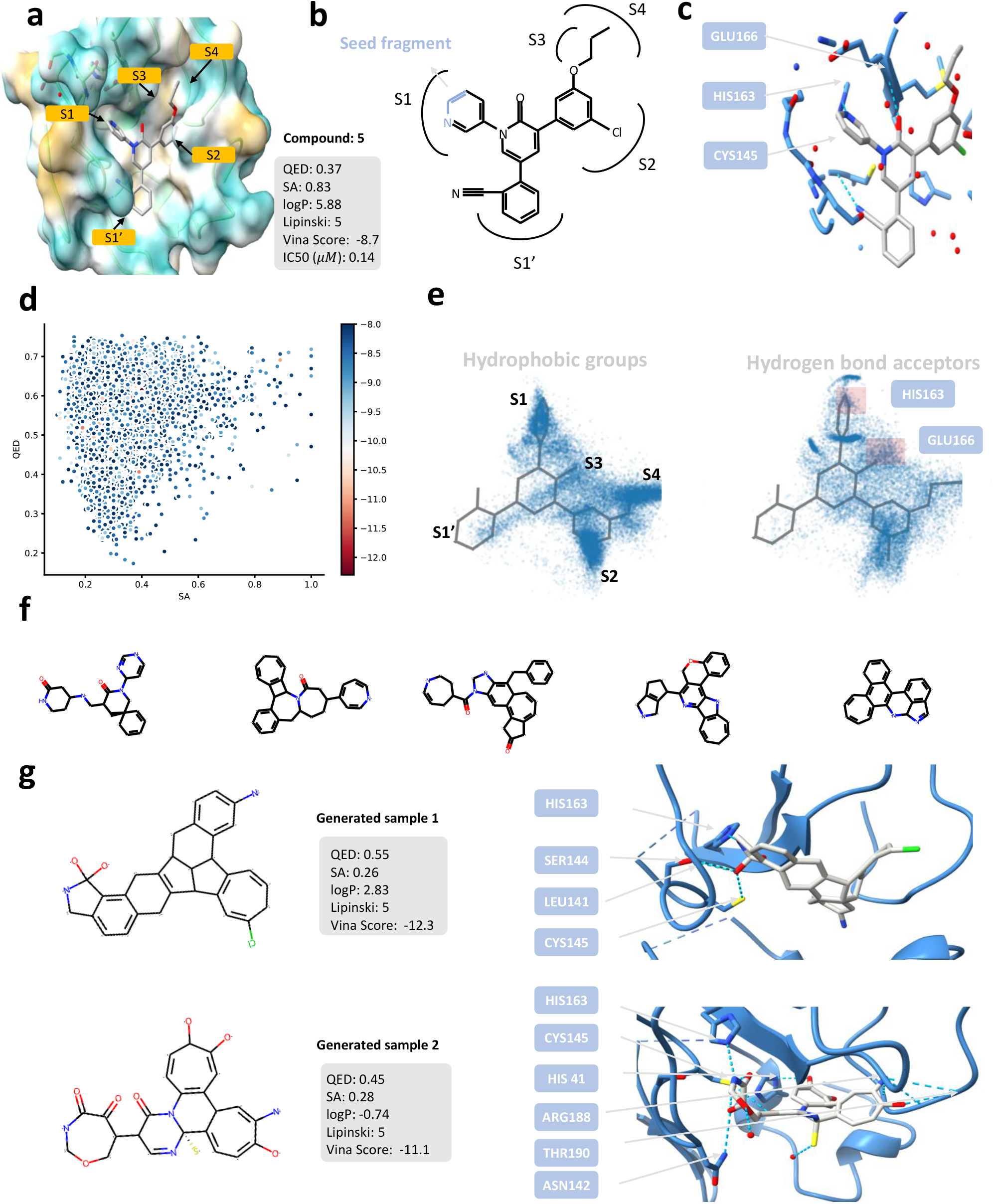
Lead optimization case of SARS-CoV-2 main protease (M^pro^). a. The complex structure of noncovalent and nonpeptidic inhibitor Compound 5 targeting M^pro^ with the pharmacochemucal properties. b. The structure of the compound 5. The blue part is the seed fragment which we utilize to generate the novel molecules. c. The hydrogen bonds between compound 5 and M^pro^. d. The Vina score, QED, and SA distribution of the generated molecules with high affinities. e. The spatial distribution of the key pharmacophore groups of generated molecules with high affinities. f. Examples of the scaffolds of generated molecules with high affinities. g Two examples of generated molecules with high affinities.

We aim to generate molecules with more novel scaffolds. Toward this end, we utilize three atoms as the seed fragment which is the blue part of Figure 5b. We generate 40000 molecules and filter out those whose vina scores are smaller than -8.0 kcal/mol. Finally, we obtain 10627 molecules with high affinities. We checked all the filtered molecules, and none of them is presented in the training set. It indicates that PMDM can still generate novel molecules binding well to the targeting proteins despite the high affinity of the reference molecule. As demonstrated in Figure 5, we plot the distribution of three key properties (QED, SA, and Vina score) of the filter molecules. As we can observe, PMDM is capable of generating molecules with good affinities while containing nice properties. Statistically, the average QED value of the molecules is 0.57, which is higher than the reference compound 5, and the maximum QED value is 0.75. For Vina score, the average value is -8.6 and the minimum value is -12.3 despite sacrificing performance in terms of synthetic accessibility which the average SA value is 0.30 and the maximum SA value is 1.0. The results demonstrate that PMDM can learn the distribution of the training data. Thus it could generate the molecules that adapt to the pocket structure and satisfy the requirement for high drug-likeness and good synthetic accessibility without inputting the desired properties as conditional information.

As we mentioned before, compound 5 contains several features contributing to its high affinity with the M^pro^. In order to investigate whether the generated molecules contain the same features, we first calculate the pharmacophore models using the software Align-It^47^. We selected the hydrophobic groups including aromatic ring(AROM), lipophilic region(LIPO), and aromatic and lipophilic(HYBL), to visualize the spatial distribution. As shown in Figure 5e, the hydrophobic groups are clustered in the S1’, S1, S2, S3, and S4, which is in accordance with the compound 5, revealing the reducing capacity of PMDM. The visual inspection of the hydrogen bond acceptors demonstrates that the interactions of HIS 163 and GLU166 are covered by the generated molecules and the position of the hydrogen bond donors aligns well with those of compound 5. Besides, there are other cluster regions which suggest that the molecules also form hydrogen bonds with the residues of the pocket.

Since we only incorporate a small fragment which only contains three atoms as the seed fragment, PMDM manages to generate molecules with more novel scaffolds. Finally, we extracted 10207 Bemis-Murcko scaffolds by RDKit from the 10627 filter molecules. Figure 5f shows examples of the novel scaffolds. The scaffolds reflect a shared commonality that all the scaffolds contain multiple rings, especially aromatic rings. The rings occupy the key binding sites (S1, S2, S3, and S4), which is key binding sites of M^pro^. Besides, we found that there exits scaffolds similar to that of the reference molecules. Specifically, the third, fourth, and fifth example scaffolds shown in Figure 5f consist of the aromatic ring connected to three rings.

The results imply that PMDM can discover the optimized structure patterns which are verified by the reference molecule. To further investigate the quality of generated molecules, we selected two compounds with improved Vina scores. We searched PubChem, ChEMBL, and DrugBank and found the two compounds are not recorded in all the datasets. Both compounds form similar interaction patterns with multiple residues of M^pro^. In addition to the hydrogen bond with residue HIS163, the compounds form hydrogen bonds with more residues to achieve higher binding affinities. Specifically, the hydroxyl group besides the seed fragment in the generated sample 1 forms three hydrogen bonds with three residues: SER144, LEU141, and CYS145. Furthermore, generated sample 1 contains Chlorine connected by an aromatic ring, which is similar to the reference compound. For the generated sample 2, the hydroxyl groups form hydrogen bonds with CYS145, HIS41, and THR190. Interestingly, the Sulfur atom also forms a hydrogen bond with the Oxygen atom. These results spotlight that PMDM can generate novel molecules highly binding to the targeted proteins. We anticipate that the web-lab experiments could be conducted to verify the inhibitory effect of the generated novel molecules on M^pro^.

## Discussion and conclusion

In this paper, we proposed a novel diffusion model, PMDM which enables 3D small-molecule ligands generation conditioned on specific target proteins in a one-shot manner by incorporating the diffusion framework. PMDM utilizes a dual equivariant encoder to handle different (global & local) molecular dynamics. To achieve protein-conditioned generation, PMDM employs the cross-attention mechanism to consider the protein semantic information by fusing the protein representation and the ligand representation in a shared high-dimension space and incorporates the whole pocket as the input of the equivariant kernel in which the protein spatial information is fixed across the neural net layers, to consider the protein structure information. With much less complexity and sampling time, it achieves substantially better or competitive performance against the SOTA methods. We anticipate that PMDM can advance the *de novo* drug optimization targeting the specific protein and accelerate future research in drug development.

## Methods

### Preliminary

Let 𝒢 = (*x, r*) denote the 3D molecular geometry where *x* = (*x*_1_, *x*_2_, …, *x*_*n*_) ∈ {0, 1} ^*n× f*^ denotes the one-hot encoded atom features, and *r* = (*r*_1_, *r*_2_,, *r*_*n*_) ∈ ℝ^*n×*3^ denotes the atom coordinates as depicted in Figure 1c. Specifically, we denote 3D ligand geometry as 𝒢 ^*L*^ = (*x*^*L*^, *r*^*L*^) and 3D protein pocket geometry as 𝒢 ^*P*^ = (*x*^*P*^, *r*^*P*^). We denote 𝒢_*t*_ for *t* = 1, …, *T* as a sequence of latent geometries where *t* indicates the index of diffusion steps.

### Background

The diffusion model^23^ is formulated as two Markov chains: *diffusion process* and *reverse process* (a.k.a denoising process). The diffusion process iteratively adds Gaussian noises to the data according to a variance preserve schedule while the reverse process gradually refines the data until it recovers the real data by eliminating the noise. The refined goal of the diffusion model is to learn the reverse process via a parameterized neural network.

The diffusion process gradually diffuses the real data distribution into a predefined noise distribution with the time setting 1 … *T*. The transformation in every time step is set as a Gaussian distribution. This whole process is then formulated as a fixed Markov chain that gradually adds Gaussian noise to the data with a variance schedule *β*_1_ … *β*_*T*_ (*β*_*t*_ ∈ (0, 1)):

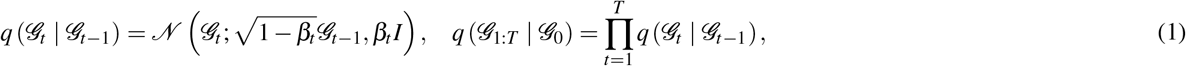

where 𝒢_*t*−1_ is mixed with the Gaussian noise to obtain 𝒢_*t*_ and *β*_*t*_ controls the extent of the mixture. By setting 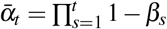, a delightful property of the diffusion process is achieved that any arbitrary time step, *t*, sampling of the data has a closed-form formulation via a reparameterization trick as:

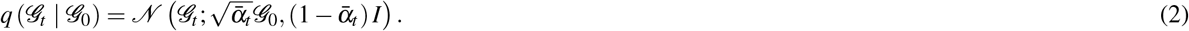

We can observe that the final distribution will be closer to a standard Gaussian distribution if the time step is large enough.

The reverse process is designed to recover the real data 𝒢_0_ from the diffused data 𝒢_*T*_ ∼ *p*(𝒢_*T*_) which is achieved by the diffusion process. The reverse process is also a Markov chain with learnable parameters which can be formulated as:

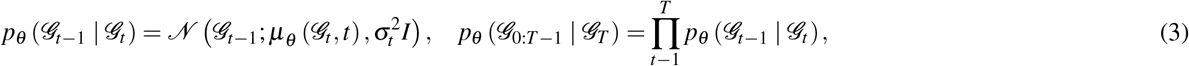

where *μ*_*θ*_ denotes the parameterized neural networks to approximate the mean, and 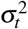 denotes user-defined variance. Specifically, we follow previous work^24^ to paramterize *μ*_*θ*_ as:

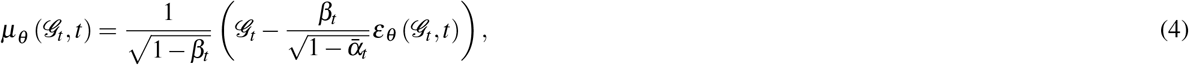

where *ε*_*θ*_ is a neural network w.r.t trainable parameters *θ*. Having formulated the reverse process, we could maximize the likelihood of the training data as our object. Since directly calculating the likelihood is intractable, we adopt the variational lower bound (VLB)^24^ to optimize.

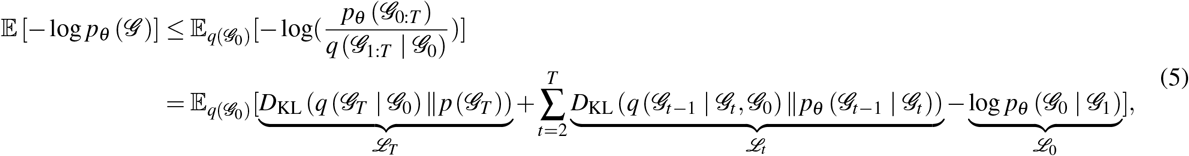

where *q*_*ϕ*_ (·) denotes a learnable variational noising encoder. The detailed derivation is left in the Appendix. ℒ_*T*_ is a constant and ℒ_0_ can be approximated by the product of the PDF of 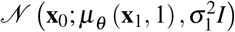 and discrete bin width. Hence, we adopt the simplified training objective as follows:

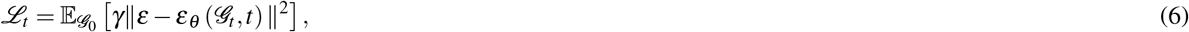

where 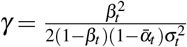 refers to a weight term. We can observe that the terminal goal of the reverse process is to learn the noised added in the diffusion process. Actually, *ε*_*t*_ can be represented as 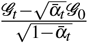 from Eq.(2) via the reparameterize trick, where 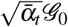 is the mean *μ* and 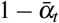 is the variance *σ* ^2^. Since the logarithmic gradient of *q* (𝒢_*t*_ | 𝒢_0_) can be formulated as 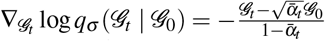, then we can obtain that 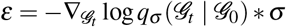. In other words, the purpose of the diffusion model is equivalent to moving the data distribution to the high-density region of the distribution led by the logarithmic gradient which initially starts from a low-density region. Therefore, the negative modified eliminated noise part −*ε*_*θ*_ *σ* is also regarded as the *(stein) score*^48^, the logarithmic density of the data point at every time step. Now we can rewrite Eq.(4) as:

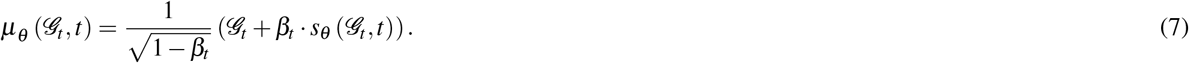

### PMDM: Pocket based Molecular Diffusion Model

In this section, we will elaborate on our proposed model PMDM: Pocket based Molecular Diffusion Model. Different from the pure diffusion model, PMDM is a conditional diffusion model instead, where the pocket protein guides the molecule generation. Thus, we attempt to model the *p*_*θ*_ (𝒢 ^*L*^ | 𝒢 ^*P*^) to obtain the distribution of the ligand binding to the pocket protein. The conditioned pocket protein semantic information is achieved by the cross-attention layer, which is effective for fusing various modalities. Specifically, we design a dual equivariant diffusion model for learning and generating the binding molecule geometry. Based on our previous model MDM^16^, we devise two equivariant kernels to simulate the local chemical bonded graph and the global distant graph. In order to ensure the relative distance between the ligand and the protein, we employ an equivariant graph neural network EGNN to handle the whole pocket which can treat the pocket geometry as the condition information. Figure 1b presents an overview of PMDM framework. Will elaborate on each component of PMDM in the following sections.

#### Conditioned protein semantic information encoder

Here, we adopt an invariant graph neural network SchNet to encode the protein semantic information first. Formally, the updates of protein node features are computed as follows:

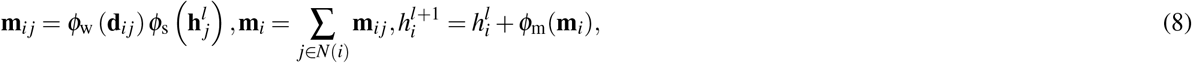

where ϕ_w_ denotes a weight network, ϕ_s_ and ϕ_m_ are multilayer perceptrons (MLPs), *d*_*i j*_ denotes the Euclidean distance between atom *i* and atom *j* of the pocket protein, and *N*(*i*) is the radius neighbourhood of atom *i*. We obtain the protein vector of the first hidden layer by a single leaner layer: *h*^0^ = Linear(*x*^*P*^). We denote the final output of the protein encoder as *h*_*P*_ for a clear description. Similarly, we also employ another SchNet to project the ligand atom feature into an intermediate representation:

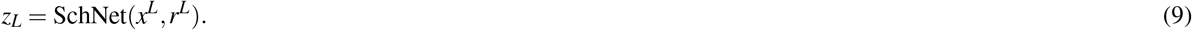

We implement the cross-attention mechanism to fuse the protein semantic information and ligand hidden information:

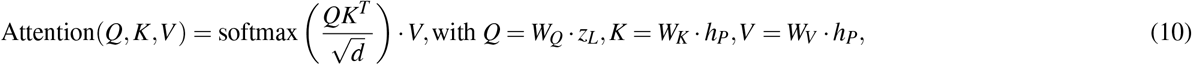

where 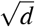 turns the attention matrix into a standard normal distribution. Specifically, the protein information is considered as the query to compute the attention score. The output of the cross-attention layer incorporates the protein semantic information as the conditioned context.

### The dual equivariant score kernels

As the molecular geometries are *invariant* to rotations and translations, we should take this property into account when devising the Markov kernels. In essence, Kohler et al.,^49^ proposed an equivariant invertible function to transform an *invariant* distribution into another *invariant* distribution. This theorem is also applied to the diffusion model^50^. If *p*(𝒢_*T*_) is *invariant* and the neural network *q*_*θ*_ which learns to parameterize *p*(𝒢_*t*−1_ 𝒢_*t*_) is *equivariant*, then the distribution *p*(𝒢_0_) is also *invariant*. Therefore, we utilize an *equivariant* Markov kernel to achieve this desired property.

### Edge construction

As we mentioned before, molecular geometries in 3D generation are represented as point clouds. Thus, we need to construct edges manually for the point clouds to feed them into the subsequent equivariant kernels. Previous works^49,18^ consider the fully connected edges to feed into the equivariant graph neural network. However, the fully connected edges connect all the atoms and treat the interatomic effects equally but regret the effects of covalent bonds. Besides, the redundant edges contain meaningless information, leading to inefficiency. Therefore, we further define the edges whose lengths are shorter than the radius *τ*_*l*_ as local edges to simulate the covalent bonds and the edges whose lengths are between *τ*_*l*_ and *τ*_*g*_ as global edges to capture the long-distance information such as van der Waals force, which is shown in Figure 1d.

Practically, we set the local radius *τ*_*l*_ as 3Å which could include almost all the chemical bonds and the global radius *τ*_*g*_ as 6Å. The one-hot encoded atom features and coordinates with the local edges and global edges are fed into the dual equivariant encoders, respectively. Specifically, the local equivariant encoder models the intramolecular force such as the real chemical bonds via local edges while the global equivariant encoder captures the interactive information among distant atoms such as van der Waals force via global edges.

### Conditioned protein spatial information

In addition to the conditioned protein semantic information, we also need to consider the conditioned protein spatial information to ensure the generated ligand can fit the pocket structure without the clash issue. Here, we combine the ligand and protein as the complete pocket as the input of the equivariant kernel. Thus, we construct the local edges and global edges for the input pocket. Specifically, we only construct the edges within the ligand and the edges within the protein to avoid cross-modal distance inference.

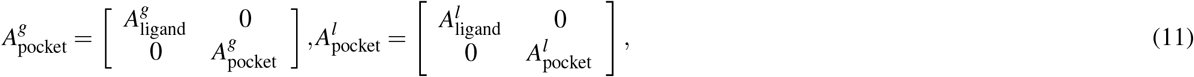

where 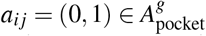 and *τ*_*l*_ *< d*_*i, j*_ ≤ *τ*_*g*_ if *a*_*i j*_ = 1, and 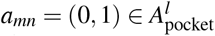 and *d*_*i, j*_ ≤ *τ*_*l*_ if *a*_*mn*_ = 1. It should be noted that we also remove the self-loop edges to eliminate replicated calculations. By constructing such separate edges, PMDM can perceive the shape of the pocket hole and ensure that the ligand can aggregate the neighborhood information independently via the message-passing process of graph neural networks. Since the pocket spatial information is treated as the condition, we keep the protein position fixed during the update of each layer of the equivariant kernel.

### Equivariant kernel

We employ E(n) Equivariant Graph Neural Networks (EGNN)^17^ to achieve the equivariant property. Here, EGNN is equivariant w.r.t the SE(3) group: EGNN(**A** 𝒢 + **b**) = **A**EGNN(𝒢) + **b** where **A** is an orthogonal rotation matrix and **b** is a translation vector. Here, we concatenate the ligand atom embeddings which already contain the protein semantic information and pocket atom features as *x*^0^ = [*z*_*L*_, *h*_*P*_], and the ligand atom coordinates and the protein coordinates as *r*_0_ = [*r*_*L*_, *r*_*P*_]. Specifically, the equivariant convolution layer takes the node embeddings **x**^*l*^ ∈ ℝ^*n×d*^, corresponding coordinate embeddings **r**^*l*^ ∈ ℝ^*n×*3^ and edge information *e*_*i j*_ as inputs at layer *l* and outputs **x**^*l*+1^ and **r**^*l*+1^. Formally, the updates of node feature and coordinate embeddings of each layer are computed as follows:

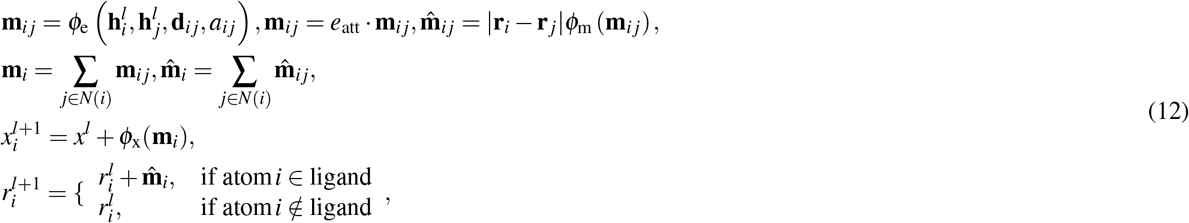

where *ϕ*_e_, *ϕ*_m_, and *ϕ*_x_ are MLPs, and *a*_*i j*_ = MLP(**d**_*i j*_) is the edge length embedding. *e*_att_ = *ϕ*_*in f*_ (**m**_*i j*_) where *ϕ*_*in f*_ : ℝ^*n×d*^ → [0, 1]^1^ is to estimate the edge value by an attention mechanism. Here, we only update the coordinates of ligands to stay the protein spatial context fixed at each layer of EGNN.

Then only the node embeddings and coordinate embeddings of the ligand part of the final layer are reserved. Finally, we add the outputs of the local equivariant kernel and the global equivariant kernel to obtain the corresponding *s*_*θ*_ :

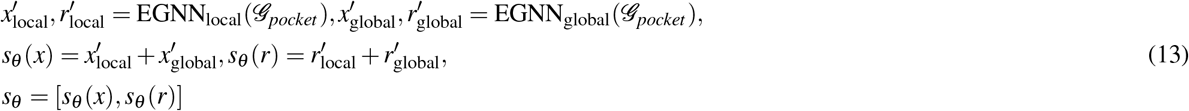

### Training

The goal of the diffusion model is to learn to reverse the diffusion process. Recall Eq.(6), we also adopt the ELBO objective of the loss function. The differences here are that we have considered the protein context information and converted the *ε*_*θ*_ to *s*_*θ*_, thus the loss function becomes:

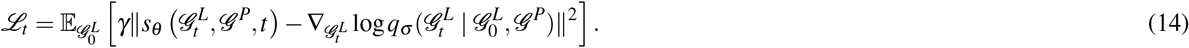

As shown in Figure 1c, PMDM sample *t* from the Uniform distribution for training every iteration. From another perspective, it ensembles *t* small models to learn the reverse process. Have achieved the equivariance of *s*_*θ*_, we also need to take this property of the coordinates of 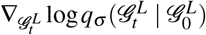 into account. Hence, we calculate 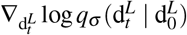 instead of 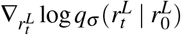 via the chain rule^51^:

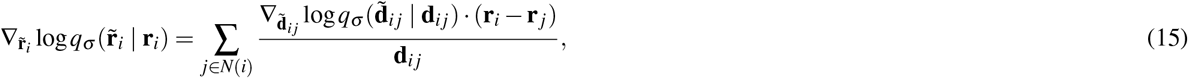

where 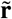 denotes the diffused atom coordinate 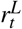 and 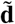 denotes the corresponding diffused distance. We approximately calculate 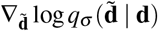 as 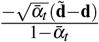

Empirically, if *γ* in Eq. (14) is ignored during the training phase, the model performs better than instead with the simplified objective. Such a simplified objective is equivalent to learning the *s*_*θ*_ in terms of the gradient of log density of data distribution by sampling the diffused molecule 𝒢_*t*_ at a stochastic time step *t*.

### Sampling from scratch

Since we have formulated the model of *sθ*, now we could calculate the *μ*_*θ*_ by Eq.(4). As presented in Figure 1a, the chaotic state 𝒢_*T*_ is sampled from *N* (0, *I*) and *μ*_*θ*_ is obtained by the dual equivariant encoder, given the target pocket protein. The next less chaotic state 𝒢_*T*−1_ is generated by 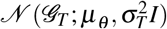. The final molecule 𝒢_0_ is generated by progressively sample 𝒢_*t*−1_ for *T* times. We adopt OpenBabel^52^ to build the chemical bonds according to the atom pairwise distances (Figure 1b). For the generic structure-based molecule generation, we adopt this sample strategy.

### Sampling from given fragment

Different from the sampling strategy from scratch which samples the molecule noise from the standard Gaussian distribution, the given fragment information 𝒢_*f*_ should be fixed as a seed start point. Here, we adopt a masked strategy to simulate the sampling process from scratch. During each iteration, the seed fragment is masked by the diffusion process according to the corresponding time step,

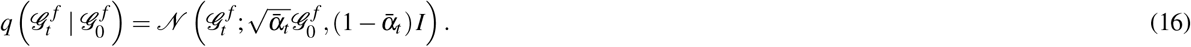

The manually diffused fragment is denoised together with the part denoised in the previous step,

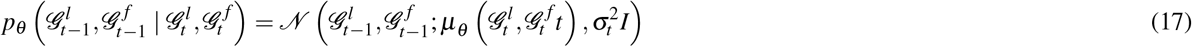

We drop the denoised fragment data 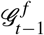 and only retain the rest of the denoised part 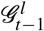 for the next iteration. Finally, we combine the fragment data and the denoised part to obtain the complete molecule by OpenBabel. For lead optimization, we adopt this sample strategy.

## Data Availability

CrossDocked dataset is available at https://bits.csb.pitt.edu/files/crossdock2020/

## Author contributions statement

Lei Huang developed and implemented the algorithms and wrote the manuscript. Tingyang Xu provided advice on the model development and case studies and revised the manuscript. Ka-Chun Wong and Hengtong Zhang conceived and supervised the project. Yang Yu provided advice on case studies and revised the manuscript. Peilin Zhao revised the manuscript.

## Notes

### Competing Interest Statement

The authors have declared no competing interest.

